# BBX16 mediates the repression of seedling photomorphogenesis downstream of the GUN1-GLK1 module during retrograde signaling

**DOI:** 10.1101/2021.11.08.467520

**Authors:** Nil Veciana, Guiomar Martín, Pablo Leivar, Elena Monte

## Abstract

- Plastid-to-nucleus retrograde signals (RS) initiated by dysfunctional chloroplasts impact photomorphogenic development. We previously showed that the transcription factor GLK1 acts downstream of the RS-regulator GUN1 in photodamaging conditions to regulate not only the well-established expression of photosynthesis-associated nuclear genes (*PhANGs*) but also to regulate seedling morphogenesis. Specifically, the GUN1/GLK1 module inhibits the light-induced PIF-repressed transcriptional network to suppress cotyledon development when chloroplast integrity is compromised, modulating the area exposed to potentially damaging high light. However, how the GUN1/GLK1 module inhibits photomorphogenesis upon chloroplast damage remained undefined.
- Here, we report the identification of *BBX16* as a novel direct target of GLK1. *BBX16* is induced and promotes photomorphogenesis in moderate light and it is repressed via GUN1/GLK1 after chloroplast damage. Additionally, we show that BBX16 represents a regulatory branching point downstream of GUN1/GLK1 in the regulation of *PhANG* expression and seedling development upon RS activation.
- The *gun1* phenotype in lincomycin and the *gun1-like* phenotype of *GLK1OX* are markedly suppressed in *gun1bbx16* and *GLK1OXbbx16*.
- This study identifies BBX16 as the first member of the BBX family involved in RS, and defines a molecular bifurcation mechanism operated by GLK1/BBX16 to optimize seedling deetiolation, and to ensure photoprotection in unfavorable light conditions.

## Introduction

To cope with their sessile condition, plants need to optimize their growth and development in response to changes in their habitat. Light is a critical environmental component necessary for photosynthesis and for the regulation of growth and development (Arsovski *et al*., 2012). Required as a primary source of energy and as an informative cue, light also represents a challenge for plant life when in excess. Plants have therefore evolved exquisite methods for light sensing and signaling to allow the appropriate adaptive response. Light of different wavelengths is perceived by different photoreceptors. Phytochromes sense red and far-red light (600-750 nm), whereas cryptochromes, phototropins, and Zeitlupes perceive blue and UVA (320-500 nm) and UVR8 senses UVB (Galvão & Fankhauser, 2015). Light perception by photoreceptors can be complemented by chloroplasts, which act as sensors of environmental changes and contribute to responses in high light (Chan *et al*., 2016).

One of the most dramatic developmental transitions in plants is deetiolation, whereby a germinating seedling experiences light for the first time (Arsovski *et al*., 2012; Gommers & Monte, 2018). When germinating in the dark, skotomorphogenic seedlings growing heterotrophically exhibit fast-growing hypocotyls, unexpanded and appressed cotyledons with etioplasts, and formation of an apical hook to protect the apical meristem from damage. In the light, deetiolated or photomorphogenic seedlings adapt their morphology to enhance light capture for photosynthesis, which involves inhibition of hypocotyl elongation, hook unfolding, stimulation of cotyledon separation and expansion, and formation of the photosynthetic apparatus and fully functional chloroplasts.

Distinct transcriptomic landscapes underlay the skoto- and photo-morphogenic programs, regulated by a suite of positive and negative acting factors (Ma *et al*., 2001; Jiao *et al*., 2005; Shi *et al*., 2018; Pham *et al*., 2018; Jing & Lin, 2020). Major positive regulators are HFR1, HY5/HYH, and LAF1 (Lau & Deng, 2012; Xu *et al*., 2015, 2016), whereas phytochrome-interacting factors (PIFs) act as major negative acting factors of photomorphogenesis (Castillon *et al*., 2007; Leivar & Quail, 2011; Leivar & Monte, 2014). PIFs (PIF1, PIF3-8) are basic helix-loop-helix (bHLH) transcription factors (Toledo-Ortiz *et al*., 2003) that bind to G-box (CACGTG) and PBE (CACATG) DNA elements in the dark to inhibit or activate the expression of light-induced or light-repressed genes, respectively (Leivar *et al*., 2009; Zhang *et al*., 2013; Pfeiffer *et al*., 2014). The quadruple mutant *pifq* lacking PIF1, PIF3, PIF4 and PIF5 displays a partial constitutively photomorphogenic phenotype in the dark, suggesting that PIFs promote skotomorphogenesis (Leivar *et al*., 2008; Shin *et al*., 2009). Upon illumination, phytochromes become active and trigger PIF inactivation and degradation through the 26S proteasome-mediated pathway, allowing seedlings to initiate light-regulated gene expression and follow a photomorphogenic program of development (Leivar *et al*., 2008, 2009; Pham *et al*., 2018). Additional transcription factors involved include the GOLDEN2-LIKE 1 (GLK1) and GLK2 (Chen *et al*., 2016) and members of the B-BOX family (BBX) (Khanna *et al*., 2009; Gangappa & Botto, 2014; Su *et al*., 2015; Song *et al*., 2020a). Whereas GLKs target genes involved in chlorophyll biosynthesis, light harvesting, and electron transport are necessary for chloroplast development (Fitter *et al*., 2002; Waters *et al*., 2008, 2009; Oh & Montgomery, 2014; Zubo *et al*., 2018), some BBX members have been described as general positive regulators of photomorphogenesis (eg BBX4/COL3, BBX11, BBX20/BZS1, BBX21/STH2, and BBX22/LZF1) (Datta *et al*., 2006, 2007, 2008; Chang *et al*., 2008; Fan *et al*., 2012; Xu *et al*., 2018; Job & Datta, 2021), and some as negative (eg BBX18/DBB1a, BBX19/DBB1b, BBX24/STO, BBX25/STH, BBX28, BBX29, BBX30, BBX31, and BBX32/EIP6) (Datta *et al*., 2006; Khanna *et al*., 2006; Indorf *et al*., 2007; Kumagai *et al*., 2008; Holtan *et al*., 2011; Wang *et al*., 2011, 2015; Gangappa *et al*., 2013; Lin *et al*., 2018; Heng *et al*., 2019b; Song *et al*., 2020b; Ravindran *et al*., 2021). In addition, the role in photomorphogenesis of BBX23/MIDA10 appears to be organ-specific (positive for hypocotyl elongation (Zhang *et al*., 2017) and negative for hook unfolding (Sentandreu *et al*., 2011). The protein stability of several of these transcription factors (e.g. HY5, LAF1, HFR1, BBX21, BBX22, and others) is directly modulated by the COP1/SPA complex acting as an E3 ubiquitin ligase, which interacts and targets them for degradation via the 26S proteasome pathway in darkness (Yi & Deng, 2005; Hoecker, 2017).

In Arabidopsis, chloroplast biogenesis during seedling deetiolation depends on the expression of chloroplast proteins encoded by the nuclear genome (~2000–3000) (Li & Chiu, 2010) (anterograde regulation), which are imported into the chloroplast following synthesis in the cytosol (Jung & Chory, 2010). In turn, chloroplasts can communicate with the nucleus through retrograde signaling (RS) to regulate nuclear gene expression according to chloroplast status (Kleine *et al*., 2009; Jarvis & López-Juez, 2014). This coordination between the nucleus and chloroplast genomes ensures optimized photosynthetic capacity and growth (Ruckle *et al*., 2007; Hills *et al*., 2015; Martín *et al*., 2016). Moderate light intensities during deetiolation induce expression of the PIF-repressed target gene *GLK1* (Martín *et al*., 2016), and GLK1 subsequently promote photosynthetic apparatus formation by directly inducing the expression of nuclear-encoded photosynthetic genes (*PhANGs*) such as those from the *LHCb* gene family (Waters *et al*., 2009). Under photodamaging conditions, however, RS is activated (Ruckle *et al*., 2007; Estavillo *et al*., 2011; Kindgren *et al*., 2012) leading to the repression of *GLK1* expression and down-regulation of *PhANGs* (Waters *et al*., 2009; Martín *et al*., 2016). The use of drugs such as lincomycin or norflurazon specifically inhibits plastid translation or carotenoid biosynthesis, respectively, and activates RS causing photobleaching and repression of *PhANG* expression (Oelmüller *et al*., 1986; Sullivan & Gray, 1999). *Genomes uncoupled* (*gun*) mutants exhibit *PhANG* derepression in response to these drugs, and have helped elucidate components of RS like tetrapyrroles such as heme, and GUN1 (Koussevitzky *et al*., 2007; Chan *et al*., 2016). Importantly, RS has been shown to impact light-regulated seedling development in high light environments to prevent photodamage, through a GUN1-mediated mechanism that is still not well defined (Ruckle *et al*., 2007; Martín *et al*., 2016). It is also currently unknown whether light regulation of seedling development and *PhANG* expression after RS activation operate through the same components.

We have previously shown that the RS and phytochrome pathways converge to antagonistically regulate the PIF-repressed light-induced transcriptional network (Martín *et al*., 2016). Our findings showed that GLK1 acts downstream of GUN1 to modulate not only *PhANG* expression but also seedling morphogenesis in photodamaging conditions. Specifically, GUN1/GLK1-mediated RS antagonize phytochrome/PIF signaling to inhibit cotyledon separation and expansion when chloroplast integrity is compromised, effectively reducing the area exposed to potentially damaging high light. How this is achieved is still unclear, but does not involve the reaccumulation of PIF proteins in these conditions (Martín *et al*., 2016), thus suggesting the participation of yet undefined components (Fig. S1). Here, we address the question of how the GUN1/GLK1 module inhibits photomorphogenesis upon chloroplast damage, and report the identification and characterization of *BBX16* as a novel GLK1 target. BBX16 promotes photomorphogenesis downstream of PIF and GLK1 in moderate light and is repressed via the GUN1/GLK1 module after chloroplast damage. Additionally, we show that BBX16 represents a regulatory branching point in the regulation of *PhANG* expression and seedling development upon RS activation.

## Materials and Methods

### Plant material and growth conditions

*Arabidopsis thaliana* wild-type and mutant seeds used in this study have been described previously. *gun1* (*gun1-201*) (Martín *et al*., 2016), *glk1* and *glk1glk2* (Fitter *et al*., 2002), *GLK1OX* and *GLK1OX-GFP* (both in *glk1glk2* background) (Waters *et al*., 2008) are in Col-0 background, whereas *col7, BBX16OX #10* and *BBX16OX #11* (here renamed as *bbx16-1, BBX16OX1* and *BBX16OX2*, respectively) (Wang *et al*., 2013b) are in Col-4 background. *BBX16OX* lines express the *BBX16* open reading frame under the control of the 35S promoter and were described to overexpress BBX16 ~250 fold (Wang *et al*., 2013b). *bbx16-1* is an insertional mutant from the GABI-Kat collection (GABI-639C04) with a T-DNA insertion in the second exon of *BBX16* (Wang *et al*., 2013b). A new second BBX16 mutant allele (named *bbx16-2*) was obtained from the SALK collection (SALK_036059), harboring a T-DNA insertion in the first exon (Fig. S2). The *gun1bbx16-1* was obtained by crossing *gun1-201* to *bbx16-1*, and WT (Col-0 x Col-4 background), *gun1* and *bbx16* siblings from the cross were selected to be used in the experiments shown in Fig. 4. *GLK1OXbbx16-1* and *GLK1OXbbx16-2* were generated by crossing *GLK1OX* to *bbx16-1* and to *bbx16-2*, respectively. The obtained mutants were selected to maintain the *glk1glk2* background in *GLK1OX*, and *GLK1OX* siblings from each cross were selected to be used as controls in the experiments shown in Fig. 4 and Fig. S4. Seeds were surface-sterilized in 20% bleach and 0.25% SDS for 10 minutes and plated on 0.5X Murashige and Skoog (MS) without sucrose, stratified at 4°C in the dark for 4 days, exposed to white light for 3 hours to induce germination, and then placed to the specific light conditions indicated in each experiment. For experiments done under continuous conditions, plates where placed under white light (5 μmol·m^-2^·s^-1^) or darkness for 3 days (unless otherwise indicated), except in experiments shown in Fig. 2d, performed using a light intensity of 20 μmol m^-2^ s^-1^, in Fig. 3f and Fig. 4b, where light was 10 μmol m^-2^ s^-^1, and in Fig. 3e, in which the light conditions consists in a combination of red (60%) and blue (40%) light, where light corresponds to 130 μmol m^-2^ s^-1^ and high light to 310 μmol m^-2^ s^-1^. In the text we refer to low light (<25 μmol·m^-2^·s^-1^), light (100-150 μmol·m^-2^·s^-1^), and high light (>300 μmol·m^-2^·s^-1^), whereas the specific ligh intensity used in each experiment is specified in the corresponding figure legend. For lincomycin treatments, media was supplemented with 0.5 mM lincomycin (Sigma L6004) (Sullivan & Gray, 1999). Primers sequences used for genotyping are provided in Table S1.

**Fig. 1.**
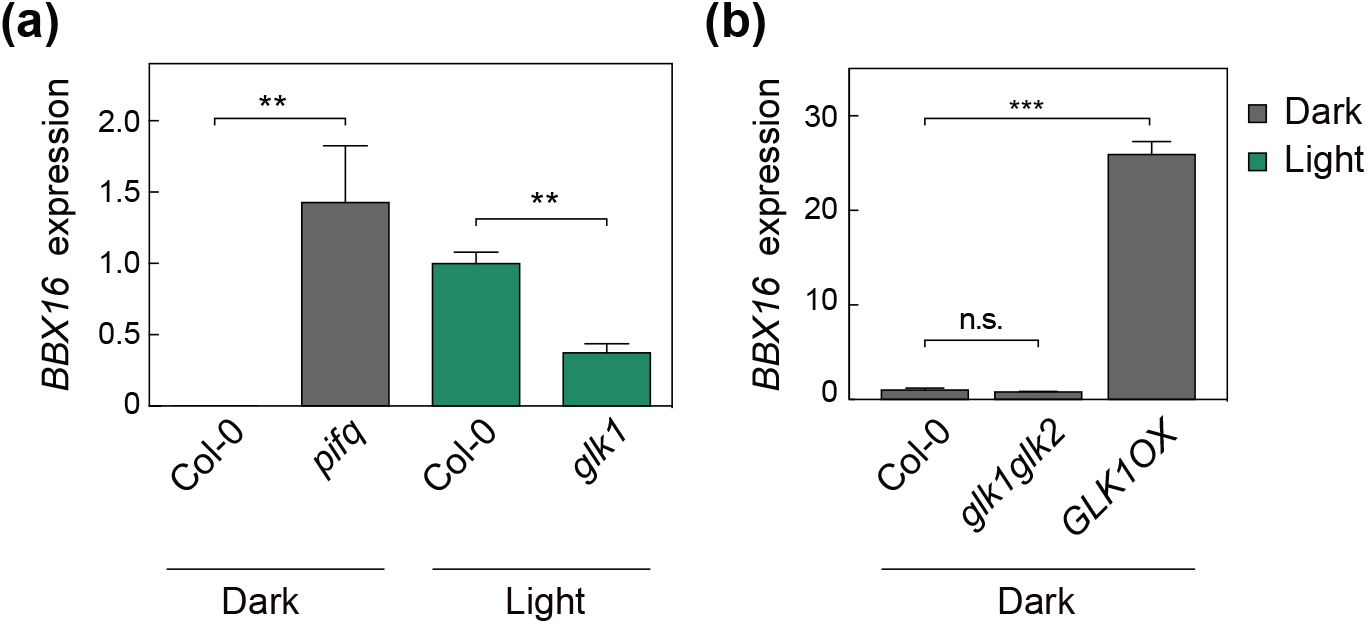
*BBX16* is a PIF-repressed gene whose expression is induced by light in a GLK1 dependent manner. **(a, b)** Transcript levels of *BBX16* analysed by qRT–PCR in (a) 3-day-old Col-0, *pifq* and *glk1* and (b) Col-0, *glk1glk2*, and *GLK1OX* seedlings grown in the dark or in continuous white light (5 μmol·m^-2^·s^-1^) as indicated. Values were normalized to *PP2A*, and expression levels are expressed relative to Col-0 light set at one. Data are the means ± SE of biological triplicates (*n*=3) and asterisks indicate statistically significant differences between each mutant and its respective WT seedlings (t-test; **P < 0.01, ***P < 0.001). n.s.: non significant.

**Fig. 2.**
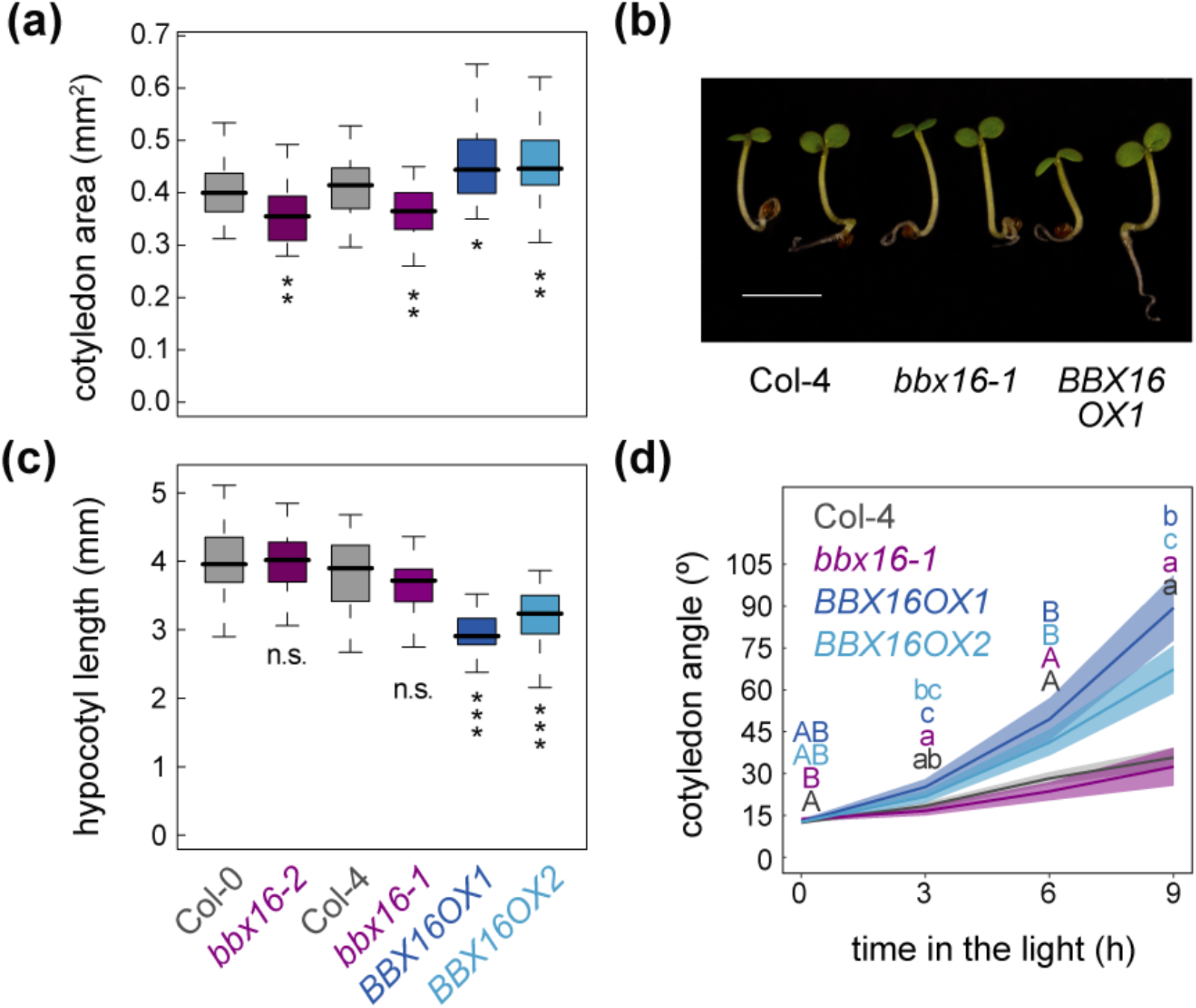
*BBX16* regulates cotyledon development during early seedling development in continuous light. **(a)** Boxplot representation of the cotyledon area of *BBX16* loss- (*bbx16*) and gain-of-function (*BBX16OX1* and *OX2*) mutants grown for three days under continuous white light (5 μmol·m^-2^·s^-1^). **(b)** Visual phenotypes of seedlings grown as detailed in (a). Bar = 2.5 mm **(c)** Boxplot representation of the hypocotyl length of seedlings grown as detailed in (a). **(d)** Quantification of the cotyledon angle of 2-day-old dark-grown WT, *bbx16* and two *BBX16* overexpressor lines transferred to white light (20 μmol·m^-2^·s^-1^) for the indicated hours (h). The thick lines and shaded areas represent the median and the 95% confidence interval of at least 60 seedlings, respectively. Letters denote the statistically significant differences between genotypes by Kruskal-Wallis test at each time point (P<0.05). (a and c) Data represent the median of at least 20 seedlings and asterisks indicate statistically significant differences between each mutant and its respective WT seedlings (Mann–Whitney test; *P < 0.05, **P < 0.01, ***P < 0.001).

**Fig. 3.**
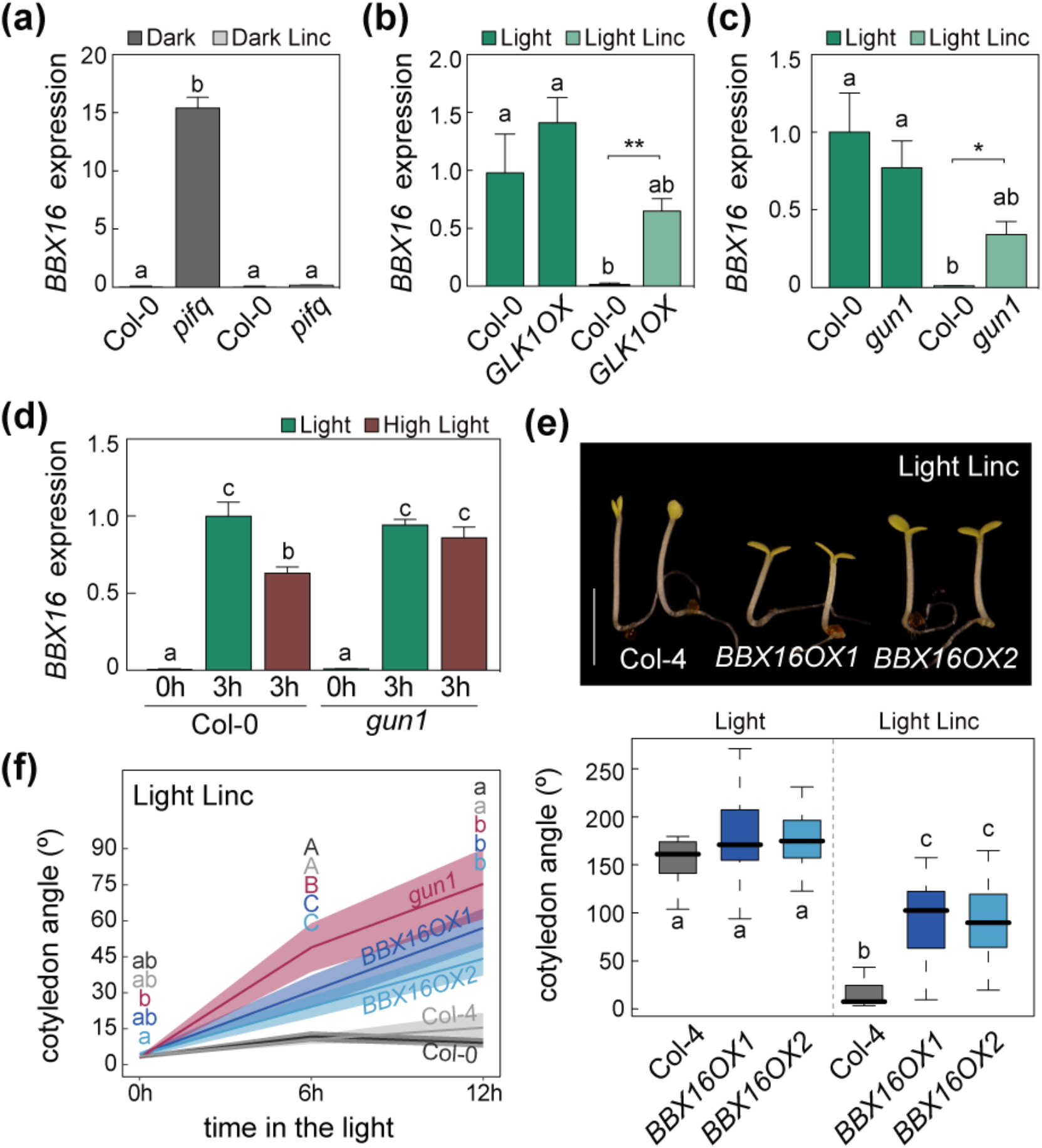
Downregulation of *BBX16* mediated by the GUN1-GLK1 module is necessary to repress cotyledon development under photo-damaging conditions. **(a)** Transcript levels of *BBX16* from RNA-sequencing of WT Col-0 and *pifq* seedlings grown for 3 days in dark in the absence or presence of lincomycin (Martín *et al*., 2016). **(b, c)** Transcript levels of *BBX16* analyzed by qRT–PCR in 3-day-old light-grown (5 μmol·m^-2^·s^-1^) Col-0 and *GLK1OX* seedlings (b), and Col-0 and *gun1* seedlings (c), in the absence or presence of lincomycin. **(d)** *BBX16* expression levels after 3 h of high light (310 μmol·m^-2^·s^-1^) compared with light (130 μmol·m^-2^·s^-1^), in WT and *gun1* mutant seedlings. (b,c,d) Values were normalized to *PP2A*, and expression levels are expressed relative to Col-0 light (b,c) or Col-0 light 3h (d), set at one. Data are the means ± SE of biological triplicates (*n*=3). (a,b,c,d) Letters denote the statistically significant differences by Tukey test (P<0.05), and asterisks in specific samples indicate statistically significant differences between each mutant and its respective WT seedlings (t-test; *P < 0.05, **P < 0.01). **(e)** Visual phenotypes (top) and cotyledon angle quantification (of at least 40 seedlings) (bottom) of WT and *BBX16OX* seedlings grown as in (b). Representative seedlings grown in presence of Lincomycin are shown in the picture. Bar = 2.5 mm. Letters denote the statistically significant differences among genotypes by Kruskal-Wallis test (P<0.05). **(f)** Quantification of the cotyledon angle of 2-day-old dark-grown WT Col-0, *gun1*, WT Col-4, *bbx16*-*1*, and two *BBX16OX* lines transferred to white light (10 μmol·m^-2^·s^-1^) for the indicated times in the presence of lincomycin. The thick lines and shaded areas represent the median and the 95% confidence interval of at least 20 seedlings, respectively. Different letters denote statistically significant differences between genotypes by Kruskal-Wallis test at each time point (P<0.05). Linc = Lincomycin.

**Fig. 4.**
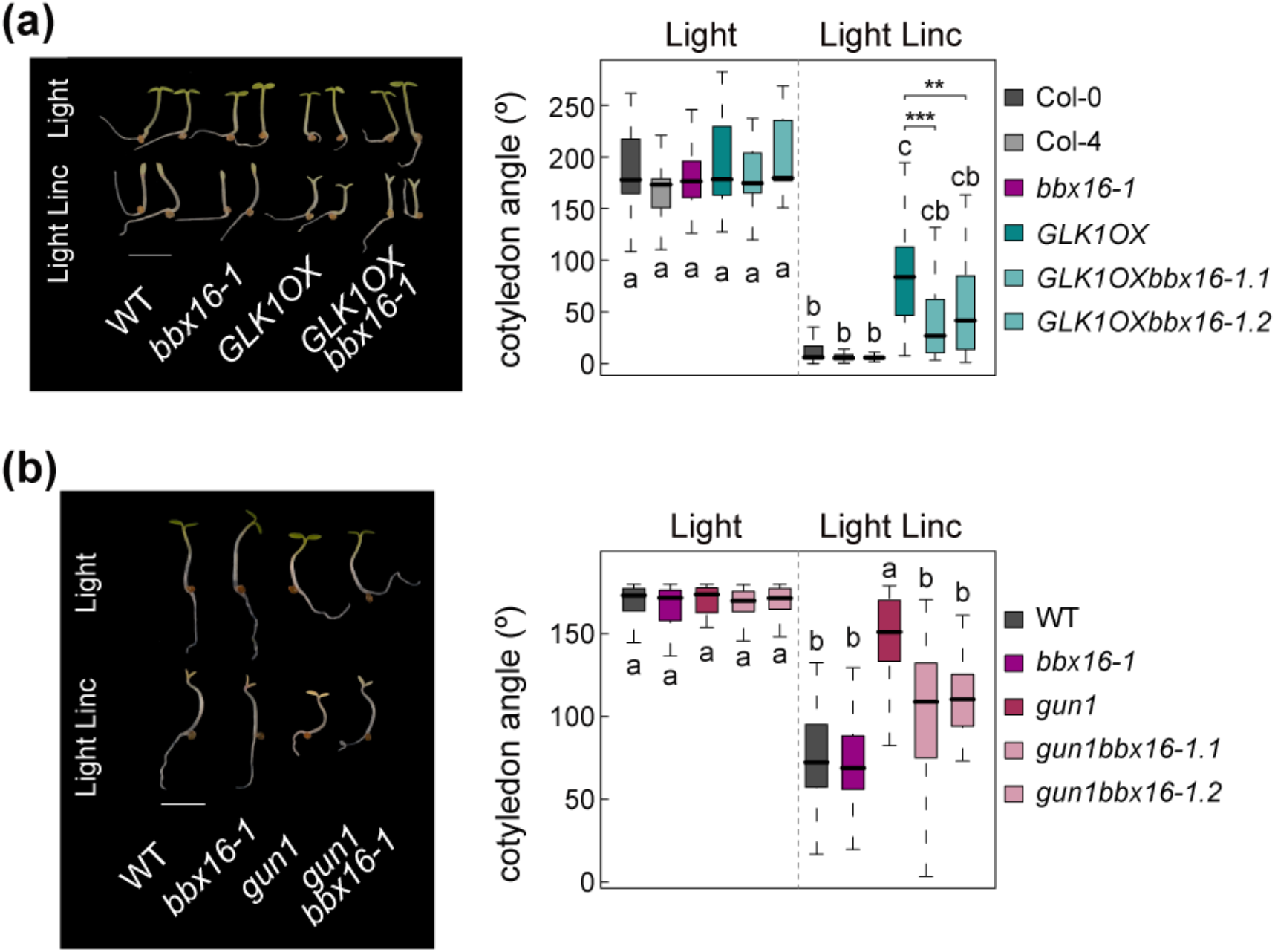
Genetic removal of *BBX16* partially suppresses the *gun1* and *GLK1OX* open cotyledon phenotype in the presence of lincomycin. **(a)** Visual phenotypes (left) and quantification of cotyledon angle (right) of 3-day old light-grown (5 μmol·m^-2^·s^-1^) Col-0, Col-4, *bbx16-1, GLK1OX*, and *GLK1OX bbx16-1* seedlings in the presence or absence of lincomycin. **(b)** Visual phenotypes (left) and quantification of cotyledon angle (right) of 2-day old dark-grown WT, *bbx16-1*, *gun1*, and *gun1bbx16*-*1* seedlings transferred to light (10 μmol·m^-2^·s^-1^) for 24 h in the presence or absence of lincomycin. (a,b) Bar = 2.5 mm. Letters denote the statistically significant differences among genotypes by Kruskal-Wallis test (P<0.05), and asterisks indicate statistically significant differences between each *GLK1OX bbx16-1* mutant and *GLK1OX* seedlings (Mann–Whitney test; **P < 0.01, ***P < 0.001). Linc = Lincomycin.

### Phenotypic measurements and statistical analysis

Hypocotyl length, cotyledon area and cotyledon aperture were measured as described (Sentandreu *et al*., 2011), by using NIH Image software (Image J, National Institutes of Health). Median was calculated from at least 20 seedlings and the experiments were repeated at least two times with similar results. Statistical analysis between genotypes was performed by the Kruskal-Wallis test (P value < 0.05), and Mann-Whitney test was used for pairwaise comparisons.

### Quantitative Reverse Transcriptase (qRT-PCR)

For qRT-PCR analysis, seedlings were grown in the dark or in white light for the indicated time. qRT-PCR was performed as described previously (Khanna *et al*., 2007) with variations. Briefly, in Figures 1a and 3b-d, 1 μg of total RNA extracted with the RNeasy Plant Mini Kit (Qiagen) was treated with DNaseI (Ambion) according to the manufacturer’s instructions. First-strand cDNA synthesis was performed using the SuperScript III reverse transcriptase (Invitrogen) and oligo dT as a primer (dT30). In Figures 1b and 6, 1μg of total RNA extracted using Maxwell^®^ RSC Plant RNA Kit (Promega) and first strand cDNA synthesis was performed using the NZY First-Strand cDNA Synthesis Kit (NZYTech). In all cases, cDNA was then treated with RNase Out (Invitrogen) before 1:20 dilution with water, and 2μl of this mix was used for real-time PCR (Light Cycler 480; Roche) using SYBR Premix Ex Taq (Roche) and primers at a 300nM concentration. Gene expression was usually measured in three independent biological replicates, and at least two technical replicates were done for each of the biological replicates. *PP2A* (*AT1G13320*) was used for normalization as described (Shin *et al*., 2007). Primers sequences used for qRT-PCR are described in Table S2.

### Chromatin Immunoprecipitation (ChIP) Assay

Chromatin immunoprecipitation (ChIP) and ChIP-qPCR assays (Fig. 4) were performed as in (Martín *et al*., 2018) using the previously described *35S::GLK1OX-GFP* line (Waters *et al*., 2008). Seedlings (3g) were vacuum-infiltrated with 1 % formaldehyde and cross-linking was quenched by vacuum infiltration with 0.125 M glycine for 5 min. Tissue was ground, and nuclei-containing cross-linked protein and DNA were purified by sequential extraction on Extraction Buffer 1 (0.4 M Sucrose, 10 mM Tris-HCl pH 8, 10 mM MgCl_2_, 5 mM ß-mercaptoethanol, 0.1 mM PMSF, 50 mM MG132, proteinase inhibitor cocktail), Buffer 2 (0.25 M Sucrose, 10 mM Tris-HCl pH 8, 10 mM MgCl_2_, 1 % Triton X-100, 5 mM ß-mercaptoethanol, 0.1 mM PMSF, 50 mM MG132, proteinase inhibitor cocktail), and Buffer 3 (1.7 M Sucrose, 10 mM Tris-HCl pH 8, 0.15 % Triton X-100, 2 mM MgCl_2_, 5mM ß-mercaptoethanol, 0.1 mM PMSF, 50 mM MG132, proteinase inhibitor cocktail). Nuclei were resuspended in nuclei lysis buffer (50 mM Tris-HCl pH8, 10 mM EDTA, 1 % SDS, 50 mM MG132, proteinase inhibitor cocktail), sonicated 10 times for 30sec each, and diluted in 10 volumes of Dilution Buffer (0.01 % SDS, 1 % Triton X-100, 1.2 mM EDTA, 16.7 mM Tris-HCl pH8, 167 mM NaCl). Overnight incubation was performed with the corresponding antibody (or with no antibody as control) at 4 °C overnight, and immunoprecipitation was performed using Dynabeads. Washes were done sequentially in Low Salt Buffer (0.1 % SDS, 1 % Triton X-100, 2 mM EDTA, 20 mM Tris-HCl pH 8, 150 mM NaCl), High Salt Buffer (0.1 % SDS, 1 % Triton X-100, 2 mM EDTA, 20 mM Tris-HCl pH 8, 500 mM NaCl), LiCl Buffer (0.25M LiCl, 1 % NP40, 1 % deoxycholic acid sodium, 1 mM EDTA, 10 mM Tris-HCl pH 8), and 1x TE. Immunocomplexes were eluted in Elution Buffer (1% SDS, 0.1M NaHCO3), de-cross-linked overnight at 65 °C in 10 mM NaCl, and then treated with proteinase K. DNA was purified using QIAGEN columns, eluted in 100 μL of QIAGEN elution buffer, and 2μL were used for qPCR (ChIP-qPCR) using *BBX16* promoter-specific primers (Table S2) spanning the regions P1 (EMP1180-P1 and EMP1182-P1) and P2 (EMP1175-P2 and EMP1176-P2) containing the predicted binding sites for GLK1 (Waters *et al*., 2009; Franco-Zorrilla *et al*., 2014), and a pair of primers inside the *BBX16* gene body as control (EMP869-P3 and EMP1177-P3) (see schematics in Fig. 5a). Three biological replicates were performed for *35S::GLK1-GFP* (Waters *et al*., 2008) incubated with and without antibody. WT controls were performed with one replicate of Col-0 seedlings with and without antibody.

**Fig. 5.**
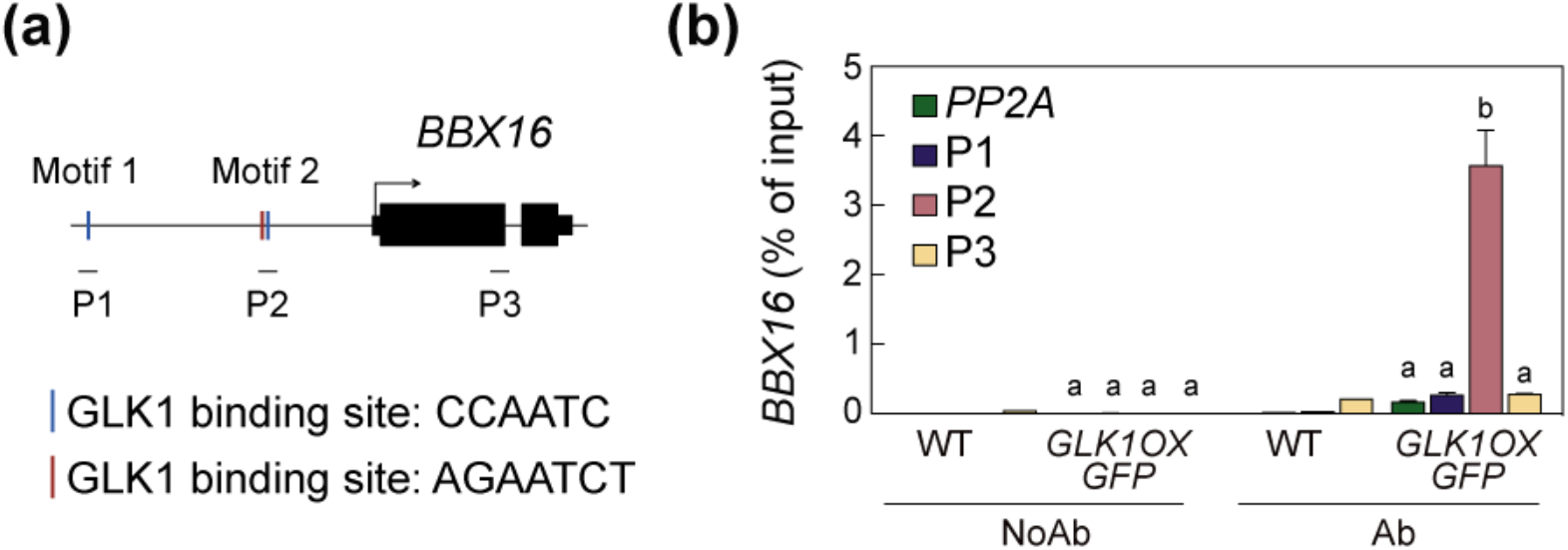
GLK1 binds to the *BBX16* promoter. **(a)** Schematic representation of the *BBX16* promoter and gene body. GLK1 binding sites (CCAATC and AGAATCT) (Waters *et al*., 2009; Franco-Zorrilla *et al*., 2014) are indicated with vertical lines in the promoter, and the regions recognized by primer pairs P1, P2 and P3 used in the ChIP-qPCR are underlined (Table S2). **(b)** GLK1 binding to the *BBX16* promoter in 3-day-old white-light (5 μmol·m^-2^·s^-1^) grown Col-0 and *GLK1OX-GFP* seedlings. Data for *GLK1OX-GFP* correspond to three independent ChIP experiments and error bars indicate the SE. Col-0 controls correspond to one biological replicate. Letters denote the statistically significant differences among *GLK1OX-GFP* samples by Tukey’s test (P<0.05). Ab, samples immunoprecipitated with antibody; No Ab, control samples immunoprecipitated without antibody.

## Results

### 1- *BBX16* is a PIF-repressed gene that is induced by light in a GLK1-dependent manner

To elucidate how the PIF/GLK1 and GUN1/GLK1 modules regulate cotyledon development under different light conditions, we aimed to identify genes downstream of GLK1 that might be involved in the regulation of photomorphogenesis. We reasoned that plausible candidates would need to meet the following criteria: (1) be a light-induced gene in a GLK1-dependent manner, and PIF repressed in dark; (2) promote cotyledon development under moderate light; (3) be a high light- and lincomycin-repressed gene via the GUN1/GLK1 module; and (4) display reduced sensitivity to RS-inducing treatments when overexpressed in seedlings, preventing RS repression of cotyledon development. Additionally, to verify the importance of the selected candidate (reresented as X), (5) genetic removal of *X* in *gun1* and *GLK1OX* mutants should suppress their phenotype in lincomycin at least partly (Fig. S1).

To begin our search, we made use of previous data describing genes directly targeted and up-regulated by GLKs (Waters *et al*., 2009). We observed that these targets (119 in total) not only included chloroplast-localized photosynthetic genes (the main focus of Waters et al. work). Significantly, we observed among them an enrichment of genes encoding for BBX transcription factors, with four of the described 32 BBX family members being present in the list of 119 genes (p-value: 2.46 e-05). Moreover, three of these BBX were members of subclass III, which is composed by four members (BBX14-BBX17). Different BBX proteins have been involved in several aspects of light-regulated development (Gangappa & Botto, 2014). In particular, BBX16/COL7 has been described to play a role in shade responses (Wang *et al*., 2013b; Zhang *et al*., 2014), and was considered a good candidate for further characterization.

To start to evaluate this candidate, *BBX16* expression was analyzed in dark- and light-grown wild-type (WT), GLK1-deficient *glk1* and *glk1glk2* (Fitter *et al*., 2002), and *GLK1*-overexpression *GLK1OX* (Waters et al., 2008) seedlings. *BBX16* was strongly up-regulated in light-grown WT seedlings compared to dark, and this induction required GLK1 (Fig. 1a). *BBX16* is a PIF-repressed gene, although not described as direct target (Pfeiffer *et al*., 2014). As such, in *pifq* etiolated seedlings, *BBX16* expression showed high levels of expression compared to WT (Fig. 1a). Interestingly, the expression of the other *BBX* in the same clade showed a similar pattern except for *BBX17* (Fig. S3), suggesting that BBX14 and BBX15 might share some function with BBX16, although the fact that *BBX16* is expressed to much higher levels (Fig. S3) might be indicative of a more important contribution. Furthermore, *GLK1* overexpression in the dark induces *BBX16* expression (Fig.1b). Together, these results indicate that during seedling establishment, *BBX16* is a PIF-repressed gene in the dark that is light-induced in a GLK1-mediated manner. Thus, the identified BBX16 met our first criterion (Fig. S1) and was considered for further genetic and molecular analyses.

### 2- BBX16 promotes cotyledon development during seedling deetiolation

Next, to evaluate the role of BBX16 during deetiolation, we analyzed the previously described *bbx16* T-DNA insertion mutant line *col7 (*referred here as *bbx16-1* for clarity), a newly characterized *bbx16-2* line (see Methods and Fig. S2), and two overexpressing *BBX16* lines (*OX1* and *OX2*) (Wang *et al*., 2013b). Under 3 days of continuous low light conditions, deficiency of *BBX16* in the *bbx16* mutants led to significantly reduced cotyledon area compared to the WT, whereas cotyledons in *BBX16-OX1 and OX2* were more expanded (Fig. 2a,b). *BBX16-OX1 and OX2* also showed slightly shorter hypocotyls (Fig. 2c). In addition, dark-grown OX lines displayed faster cotyledon aperture compared to WT after light exposure (Fig. 2d). Together, these results indicate that BBX16 contributes to the promotion of early photomorphogenesis with a role in cotyledon development (and therefore fulfills the second criterion, Fig. S1), and a possible minor contribution to the inhibition of hypocotyl elongation.

### 3- Under photo-damaging conditions, inhibition of cotyledon separation involves GUN1-mediated repression of *BBX16*

Next, *BBX16* expression was analyzed under conditions where chloroplast integrity is compromised by lincomycin treatment, an inhibitor of chloroplast translation that specifically damages the chloroplast under both dark and light conditions (Sullivan & Gray, 1999). When chloroplast is perturbed, activation of RS induces down-regulation of *GLK1* expression in a GUN1-mediated manner impacting cotyledon development (Martín *et al*., 2016). We hypothesized that under these conditions, repression of *GLK1* should also result in the repression of *BBX16* expression as a downstream effector of GLK1 (criterion 3, Fig. S1). Notably, lincomycin treatment prevented de-repression of *BBX16* in dark-grown *pifq* (Fig. 3a). Moreover, the light-induced expression of *BBX16* shown in Fig. 1 was strongly inhibited in response to lincomycin in WT seedlings (Figs. 3b, 3c), similarly to the reported inhibition of *PhANGs* and *GLK1* expression (Martín *et al*., 2016). Importantly, the inhibition of *BBX16* expression in lincomycin was only partial in *GLK1OX* (Fig. 3b), similar to *gun1* mutant (Fig. 3c).

The biological relevance of these findings using lincomycin was assessed by testing *BBX16* expression under chloroplast photo-damaging conditions. Induction of *BBX16* in high light in WT was reduced compared to normal light (Fig. 3d), suggesting that high-light damage partially inhibits *BBX16* induction, in agreement with recent transcriptomic data obtained under high light stress (Huang *et al*., 2019). This effect was not observed in GUN1-deficient mutants (Fig. 3d), indicating that this repression is mediated by GUN1. These results are in accordance with previously observed inhibition of *GLK1* under similar conditions (Martín *et al*., 2016) and suggest that the light induction of *BBX16* downstream of GLK1 is repressed in conditions where RS is active and inhibits GLK1 function.

Next, we tested whether the transcriptional repression of *BBX16* in response to RS might contribute to the inhibition of seedling deetiolation upon chloroplast damage previously observed (Martín *et al*., 2016). Indeed, *BBX16OX* lines grown for three days in plates containing lincomycin under light were less sensitive to lincomycin and were able to deetiolate, showing a cotyledon aperture that was similar to WT seedlings without lincomycin (Fig. 3e). Likewise, in a deetiolation experiment using 2-day old dark-grown seedlings transferred to light in the presence of lincomycin, *BBX16OX* lines showed reduced sensitivity to lincomycin like *gun1*, and displayed higher cotyledon angles compared to WT (Fig. 3f). These results indicate that BBX16 also fulfills criteria 3 and 4 (Fig. S1), and provide strong support that RS-imposed GUN1/GLK1-mediated repression of *BBX16* is necessary for the inhibition of cotyledon development under conditions where chloroplast is damaged.

Importantly, to provide conclusive support for this pathway, we next tested the genetic interactions between GLK1, GUN1 and BBX16 (criterion 5, Fig. S1). Genetic removal of *BBX16* in *GLK1OXbbx16* and *gun1bbx16-1* mutants allowed us to determine the contribution of the endogenous *BBX16* to the cotyledon phenotypes of *GLK1OX* and *gun1* in lincomycin (Fig. 4 and S4). Remarkably, the *gun1-like* phenotype of *GLK1OX* in lincomycin was clearly suppressed in *GLK1OXbbx16*, both in *GLK1OXbbx16-1* (Fig. 4a) and *GLK1OXbbx16-2* alleles (Fig. S4). Likewise, the *gun1bbx16-1* double mutant showed strong suppression of the open cotyledon phenotype of *gun1* (Fig. 4b). Together, we conclude that BBX16 is a promoter of cotyledon photomorphogenesis in moderate light that is targeted by the GUN1/GLK1 module under high light conditions to protect the seedling by reducing the exposed cotyledon surface.

### 4- GLK1 associates with the promoter of *BBX16*

To further understand the mechanism by which the light environment impacts development through the GLK1 regulation of *BBX16* expression, we aimed to test whether *BBX16* is a direct downstream target of GLK1 during deetiolation. Interestingly, analysis of the promoter region of *BBX16* revealed two CCAATC motifs, described as putative GLK1 binding sequences by Waters *et al*. (2009) based on the enrichment in the promoter regions of GLK1 targets. These two motifs are 2,101 bp (Motif 1) and 767 bp (Motif 2) upstream of the transcriptional start site (TSS) (Fig. 5a). Chromatin immunoprecipitation (ChIP) followed by qPCR in light-grown seedlings expressing GLK1-GFP (Waters *et al*., 2008) detected strong specific binding of GLK1 to the *BBX16* promoter specifically in the region that spans Motif 2 (P2), whereas no binding was detected to the region containing Motif 1 (P1) or a control sector within the gene body (P3) (Fig. 5b). This result indicates that *BBX16* is indeed a direct target of GLK1 during seedling deetiolation. Interestingly, we observed that the region spanning Motif 2 also contained an AGATTCT sequence in the reverse strand, identified as a potential GLK1 binding site by using protein-binding microarrays (Franco-Zorrilla *et al*., 2014). It is currently unknown whether the two binding elements in the region spanning Motif 2 are necessary for GLK1 association to the *BBX16* promoter.

### 5- BBX16 mediates regulation of only some GLK1-regulated *PhANG* genes

GLKs are key regulators of *PhANGs* (Waters *et al*., 2009; Zubo *et al*., 2018). To test whether BBX16 participates in the downregulation of *PhANG* expression in response to retrograde signals, we next studied expression of the described RS-regulated *PhANGs LCHB1.4*, *LHCB.2.2*, *CA1*, *RBCS1A*, and *RBCS3B* (Waters *et al*., 2009), in low light-grown WT, *bbx16*, *BBX16OX*, *gun1*, *GLK1OX*, and *GLK1OXbbx16-1* seedlings. In the absence of lincomycin, *LCHB1.4* and *LHCB.2.2* expression was similar to WT in all lines tested except in *GLK1OX*, where expression of both genes was upregulated as described (Waters *et al*., 2009), and in *BBX16-OX*, where *LHCB.2.2* expression was ~2-fold higher compared to WT (Fig. 6). In response to lincomycin, expression in *gun1* and *GLK1OX* lines was derepressed in accordance to Waters *et al*. (2009), whereas expression in *BBX16-OX* seedlings was similar to WT (Fig. 6). In clear contrast, expression of *CA1, RBCS1A*, and *RBCS3B* was similar to WT in all lines in the absence of lincomycin, but interestingly their expression in *BBX16OX* in the presence of lincomycin was derepressed compared to WT, similarly to *gun1* (Fig.6). Together, these results can be interpreted to suggest that BBX16 does not mediate the regulation of the *LCHB1.4* and *LHCB.2.2* upon chloroplast damage, whereas BBX16OX exhibits a *gun*-like phenotype for some *PhANGs* such as *CA1*, *RBCS1A*, and *RBCS3B*. This difference may be indicative of branching in signaling downstream of GLK1, whereby GLK1-mediated regulation of some *PhANGs* might be indirect through transcriptional regulation of *BBX16* and possibly other factors. Indeed, whereas *LCHB1.4* and *LHCB.2.2* were described as GLK1 primary targets, *CA1*, *RBCS1A*, and *RBCS3B* failed to meet the criteria to be considered in this group (Waters *et al*., 2009). Importantly, *CA1*, *RBCS1A*, and *RBCS3B* transcript levels in lincomycin were similar in *GLK1OX* and *GLK1OXbbx16* (Fig. 6). This was in contrast to the clear suppression of the *GLK1OX* cotyledon phenotype in *GLK1OXbbx16* shown above (Fig. 4), suggesting that for *PhANG* expression the contribution of endogenous BBX16 under these conditions might be relatively small.

**Fig. 6.**
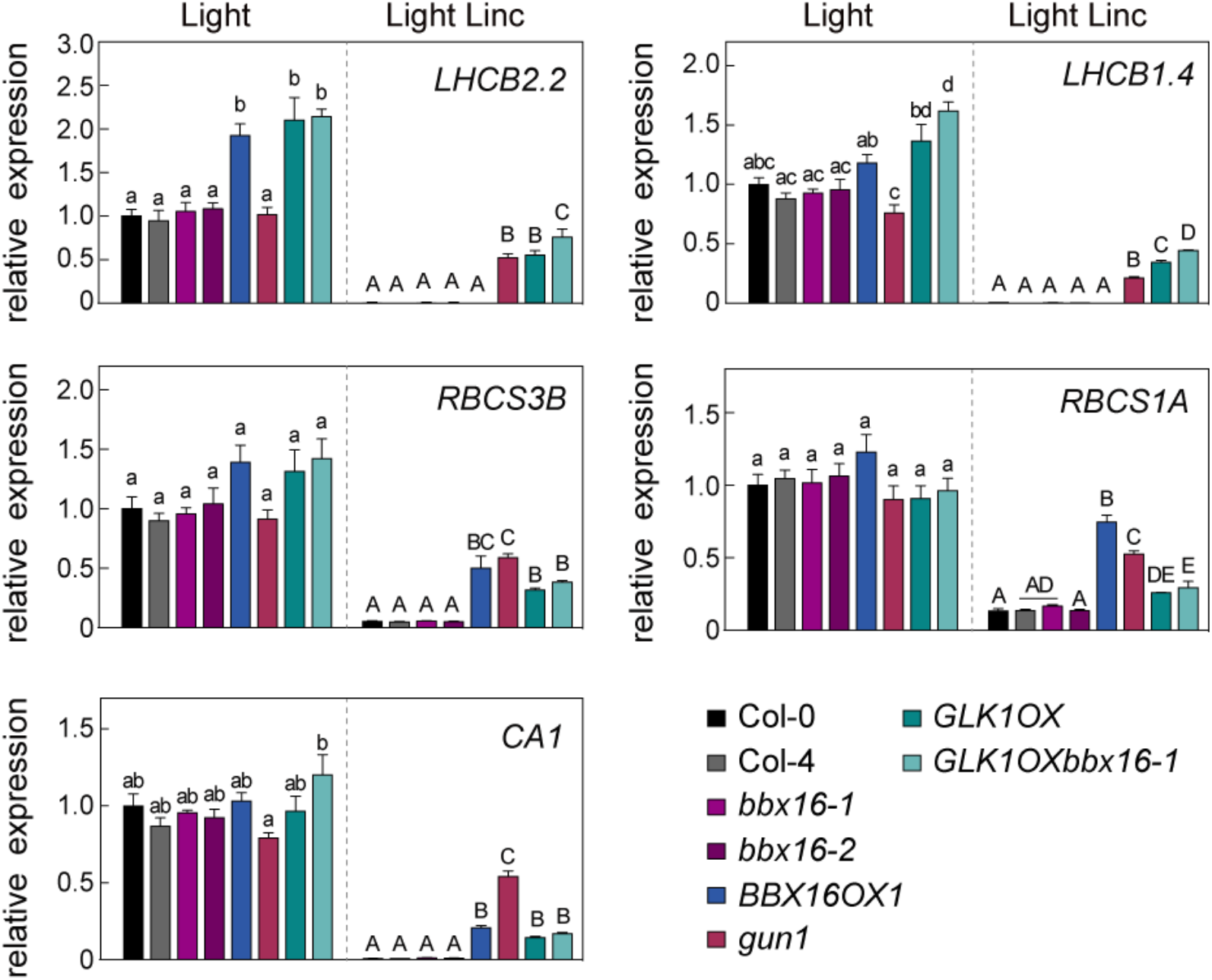
BBX16 regulation of *PhANG* genes in response to lincomycin. Expression of *LHCB2.2*, *LHCB1.4*, *RBCS3B*, *RBCS1A*, and *CA1* was analyzed by quantitative RT-PCR in WT, *bbx16*, *BBX16OX*, *gun1*, *GLK1OX*, and *GLK1OXbbx16* seedlings grown for 3 days (3d) in white light (5 μmol·m^-2^·s^-1^) in the absence or presence of lincomycin. Expression levels relative to Col-0 light are shown. Data are the means ± SE of biological triplicates. Letters denote the statistically significant differences among genotypes by Tukey’s test at each condition (P<0.05). Linc = Lincomycin.

## Discussion

The establishment of young seedlings after germination is a highly vulnerable process regulated by a myriad of factors, light being one of the most important (Gommers & Monte, 2018). Light induces transcriptional changes of hundreds of genes involved in deetiolation (Ma *et al*., 2001), many of them directly regulated by the phytochrome/PIF system, including *GLK1* (Leivar *et al*., 2009; Pfeiffer *et al*., 2014). However, because too much light is detrimental for chloroplast function and can hinder establishment, seedlings in potentially photodamaging light initiate retrograde signaling (RS) and inhibit deetiolation (Ruckle *et al*., 2007; Martín *et al*., 2016). This process is mediated by the nuclear-encoded chloroplast-localized PRR protein GUN1, which accumulates preferentially during early stages of chloroplast biogenesis and under retrograde signaling conditions (Wu *et al*., 2018), through a process that is not yet well understood but may require physically interaction with a large number of proteins (Pesaresi & Kim, 2019; Jiang & Dehesh, 2021; Wu & Bock, 2021) involved in plastid translation machinery(Tadini *et al*., 2016; Marino *et al*., 2019), tetrapyrrole biosynthesis (Shimizu *et al*., 2019), RNA editing (Zhao *et al*., 2019), and plastidial import (Khanna *et al*., 2009; Wu *et al*., 2019; Tadini *et al*., 2020). Given all these putative interactions, GUN1 has been proposed to act as scaffold protein that promotes protein complex formation (Colombo *et al*., 2016), and may allow GUN1 to function as integrator of signals from several retrograde signaling pathways. Downstream of GUN1, the nuclear localized GLKs directly regulate *PhANG* expression to inhibit chloroplast development (Waters *et al*., 2009). The GUN1/GLK1 module has also been shown to be central in the regulation of seedling morphology, although how this takes place was unknown (Martín *et al*., 2016). Here, we show that GLK1 directly induces *BBX16* to promote cotyledon development during seedling deetiolation in light conditions sustaining normal photosynthetic activity. In contrast, activation of RS under high light prevents *BBX16* upregulation through GUN1-mediated repression of *GLK1*, and this keeps the cotyledons underdeveloped to reduce the photosynthetic tissues exposed to light. Therefore, the identification of *BBX16* as a direct target of GLK1 in the regulation of photomorphogenesis defines a new molecular mechanism to optimize development during seedling deetiolation and to ensure photoprotection of the organism in unfavorable light conditions (Fig. 7).

**Fig. 7.**
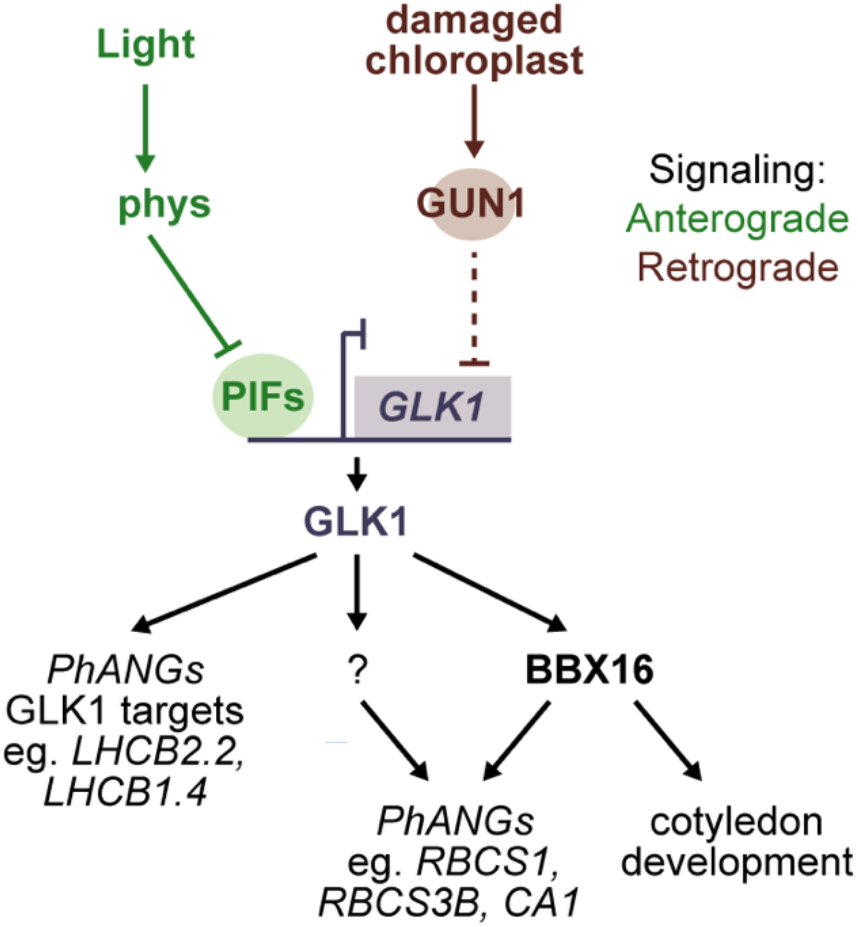
The GUN1-GLK1 module regulates *BBX16* expression during retrograde signaling. Downstream branching of GLK1 signaling directly induces two independent transcriptional pathways to regulate expression of (1) photosynthesis-associated nuclear genes (*PhANGs*) such as *LHCB2.2* and *LHCB1.4*, and (2) *BBX16* to implement cotyledon development, and indirect regulation of *PhANGs* such as *CA1*, *RBCS1A*, and *RBCS3B*, possibly with involvement of other factors (denoted as ?). In the dark, PIFs bind to the *GLK1* promoter to directly repress *GLK1* expression. In response to normal light, activated phytochromes (phys) release PIF repression on *GLK1* promoter, which triggers *GLK1* transcription. If chloroplast integrity is disrupted by lincomycin or high light, retrograde signals emitted by dysfunctional chloroplast induce GUN1-mediated repression of *GLK1* expression by a yet unknown mechanism, preventing *BBX16* and *PhANGs* transcription to block the progression of photomorphogenesis.

### BBX16 defines a signal branching hub in chloroplast-to-nucleus retrograde signaling downstream of the GUN1/GLK1 module

Our finding that GLK1 targets *BBX16* to regulate cotyledon development and to possibly regulate some *PhANGs* indirectly, whereas other *PhANGs* are directly regulated by GLK1, establishes a branching point in the regulation of seedling morphology downstream of the GUN1/GLK1 module, and indicates that the signal that GLK1 relays diversifies to specifically regulate different processes central to seedling deetiolation. Signaling network branching is common in all organisms and contributes to establishing complex responses to a given unique stimulus (Purvis et al., 2008). Interestingly, signal branching was previously described downstream of the PIFs to regulate different organ-specific pathways during seedling deetiolation (Sentandreu *et al*., 2011), where the BBX protein BBX23/MIDA10 was shown to predominantly regulate hook unfolding. Here, whereas direct GLK1 targeting of some *PhANG* genes might allow for fast regulation of chloroplast protection to e.g. fluctuations in light conditions, branching of the signal to repress *BBX16* and its target effectors would entail a slower response to arrest cotyledon development only in more sustained high light conditions, a possibility that needs further investigation.

### BBX16 is the first described BBX protein involved in Retrograde Signaling

Our finding that BBX16 is a downstream target of the GUN1/GLK1 module in RS regulated development identifies the first BBX protein involved in the response to chloroplast photodamage. This adds to previously described members of the BBX family with regulatory roles in stress-induced signaling pathways, such as BBX24/STO in responses to salt (Nagaoka & Takano, 2003), BBX18 and BBX23 to heat (Wang *et al*., 2013a; Ding *et al*., 2018), or BBX7 and BBX8 to cold stress (Li *et al*., 2021). In addition, altered expression levels of *BBX19* were found in *ceh1*, a mutant with high levels of the MEcPP retrograde signal (Xiao *et al*., 2012), although the significance is still unclear (Wang *et al*., 2014). Interestingly, a recent bioinformatic analysis of the *BBX* family identified that the promoter region of *BBX16* contains cis elements predicted to be abscisic acid, low temperature and drought responsive (Lyu *et al*., 2020), which could indicate a role for BBX16 in the cross-talk between different stress pathways.

The BBX family in *Arabidopsis thaliana* consists of 32 proteins arranged into five structural groups (I-V) based on the number of B-Box motifs (one or two) and the presence or absence of a CCT domain and a VP motif (Robson *et al*., 2001; Kumagai *et al*., 2008; Khanna *et al*., 2009; Gangappa & Botto, 2014). BBX16/COL7 belongs to the Class III clade, the least characterized of the BBX groups, together with BBX14/COL6, BBX15/COL16 and BBX17/COL8, defined by having only one B-Box motif (B-Box 1) in combination with a CCT domain. The expression patterns shown in Fig. S3 indicate that *BBX14* and *BBX15* respond similarly to *BBX16*. Because functional redundancy is common among members of the same clade within transcription factor families (Soy *et al*., 2014; Pfeiffer *et al*., 2014; Zhang *et al*., 2017; Leivar *et al*., 2020; Martín *et al*., 2020) this leads us to speculate that BBX14 and BBX15 might share some functional aspects with BBX16. Redundancy within this clade would imply that the *bbx16* mutant still retains functionality and, accordingly, we detected more prominent cotyledon phenotypes in *BBX16-OX* compared to *bbx16*. Future genetic characterization of single and high order mutant combinations in *bbx14*, *bbx15*, and *bbx16* will shed light on possible functional redundancy and address whether BBX14 and BBX15 might also play a regulatory role in response to chloroplast damage. Interestingly, a recent transcriptomic study identified Class III clade as potential players in response to high light (Huang *et al*., 2019). Of future interest will be as well to explore whether the BBX family of transcription factors has functionally evolved and diverged to specialize only Class III in RS regulation, or whether BBX factors from other clades might also be involved.

### The domain-function structure of BBX16, a promoter of photomorphogenesis

The domain structure of BBX proteins has important functional implications. B-Box domains have been involved in protein-protein interactions and transcriptional regulation, whereas the CCT harbors a nuclear localization signal (NLS) to mediate nuclear protein transport (Robson *et al*., 2001), and has also been shown to participate in the association to DNA (Ben-Naim *et al*., 2006; Tiwari *et al*., 2010). CCT-containing BBX proteins include CONSTANTS (BBX1/CO), one of the best-studied BBX proteins and the founder of the family. In CO, CCT is required to interact with the E3 ubiquitin ligases COP1 and SPA proteins (Laubinger *et al*., 2006; Jang *et al*., 2008), whereas the B-box1 domain mediates interaction with BBX19 (Wang *et al*., 2014). In the regulation of seedling photomorphogenesis, a number of BBX proteins are related to the COP1/SPA-HY5 regulatory hub (Gangappa & Botto, 2014; Song *et al*., 2020a; Xu, 2020). Several of these BBX proteins interact with COP1 and are regulated in a COP1-dependent manner, and/or regulate HY5 transcription, stability, or activity (Datta *et al*., 2006; Chang *et al*., 2011; Holtan *et al*., 2011; Jiang *et al*., 2012; Gangappa *et al*., 2013; Huang *et al*., 2014; Xu *et al*., 2016; Wei *et al*., 2016; Zhang *et al*., 2017; Job *et al*., 2018; Lin *et al*., 2018; Ding *et al*., 2018; Bursch *et al*., 2020). Furthermore, BBX4 has been shown to interact with PIF3 and repress its activity in red light (Heng *et al*., 2019a), whereas BBX18 and BBX23 have been shown to interact with ELF3 and regulate thermomorphogenesis in Arabidopsis (Ding *et al*., 2018). Whether BBX16 is regulated by the COP/SPA system, and whether BBX16 regulation of cotyledon development downstream of the GUN1/GLK1 module involves HY5 or other interacting proteins, are matters that await future research. Interestingly, the CCT domain of BBX16/COL7 has been shown to mediate binding to the promoter of the auxin biosynthesis repressor *SUR2* in the regulation of plant architecture under shade conditions in Arabidopsis adult plants (Zhang *et al*., 2014). In addition, other BBX factors like BBX20 and BBX32 have been shown to regulate photomorphogenesis through mediating brassinosteroid and strigolactone homeostasis (Wei *et al*., 2016; Ravindran *et al*., 2021). Because auxin and other hormones are well known key regulators of photomorphogenesis, and integration of retrograde and hormonal signaling is essential in the adaptation to a myriad of stresses (Jiang & Dehesh, 2021), it will be of interest in the future to explore a connection of BBX16-mediated retrograde signaling with key regulatory genes in diverse hormone pathways that could impact cotyledon development.

To conclude, this study supports a model whereby *BBX16* is directly targeted by GLK1 to induce cotyledon photomorphogenesis under light conditions favorable for seedling deetiolation. In contrast, when GUN1-mediated RS is activated, the inhibition of *GLK1*, *BBX16* and *PhANG* expression limits cotyledon and chloroplast development to minimize light damage and optimize photoprotection. The importance of this response is illustrated by studies with *gun1* seedlings exposed to high light, which exhibit more photobleached areas in their cotyledons compared to the WT controls (Ruckle *et al*., 2007). This adaptive mechanism would protect an etiolated seedling, which is extremely vulnerable, emerging into excess light such as that in a hot sunny day. This could take place transiently during establishment, allowing the seedling to prevent damage and wait safely for the light to become less strong due to shading or the natural shift in the position of the sun.

## Acknowledgements

We are grateful to Bin Liu (College of Life Sciences, Hunan University, China) for generously sharing the *bbx16*-*1/col7* and *BBX16*-*OX* lines, to Jane Langdale (University of Oxford) for *GLK1* lines, and to ABRC/NASC for *bbx16-2* seeds. This work was supported by grants from FEDER / Ministerio de Ciencia, Innovación y Universidades - Agencia Estatal de Investigación (Project References BIO2015-68460-P and PGC2018-099987-B-I00) and from the CERCA Programme / Generalitat de Catalunya (Project Reference 2017SGR-718) to E.M. We acknowledge financial support from the Spanish Ministry of Economy and Competitiveness, through the “Severo Ochoa Programme for Centres of Excellence in R&D” 2016-2019 (SEV-2015-0533)”.

## Author Contributions

EM, PL, GM and NV conceived the project and planned the experiments. GM and NV performed experiments and analyzed the data. All authors wrote the manuscript. NV and GM contributed equally.

## Data Availability Statement

The data that support the findings of this study are available from the corresponding author upon reasonable request.

## Supporting Information

**Fig. S1.** Model depicting the criteria we followed to identify putative regulators of cotyledon development downstream of GLK1.

**Fig. S2.** Molecular characterization of *bbx16-2*.

**Fig. S3.** *BBX14, BBX15* and *BBX16* are similarly regulated by PIFs and GLK1 in dark and light.

**Table S1.** List of primers used for genotyping.

**Table S2.** List of primers used for qRT-PCR.

