## Supplemental Material for "BBX16 mediates the repression of seedling photomorphogenesis downstream of the GUN1-GLK1 module during retrograde signaling"

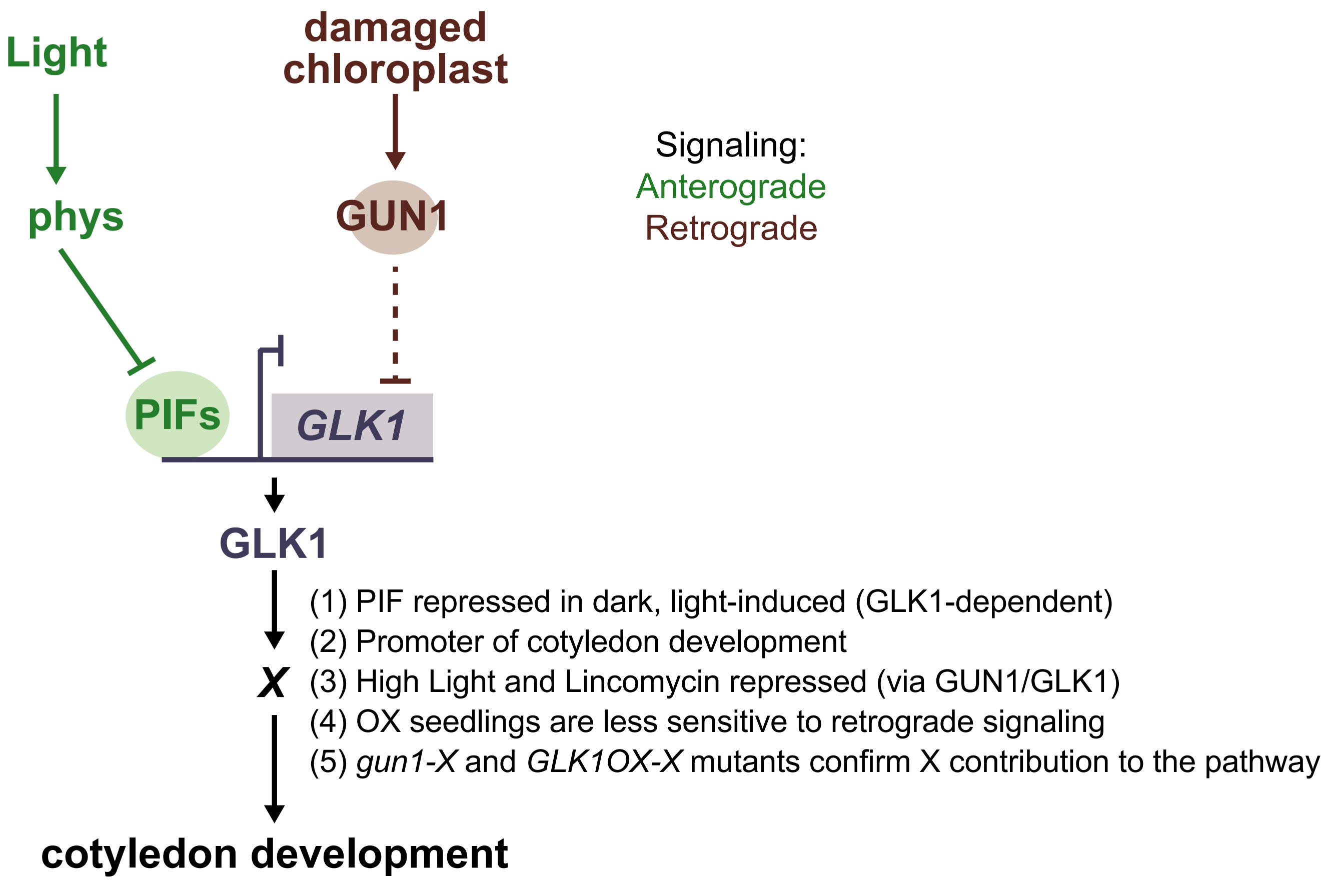


**Fig. S1. Model depicting the criteria we followed to identify putative regulators of cotyledon development downstream of GLK1.**
Based on our previous results (Martín et al., 2016), we considered that *X* should be a gene repressed transcriptionally by PIFs in the dark, and activated by light in a GLK1-dependent manner. Under normal light conditions, X promotes cotyledon development. Moreover, gene *X* is downregulated by the GUN1/GLK1 module in response to activation of retrograde signaling by high light or lincomycin. Consequently, we expect that seedlings overexpressing *X* will be less sensitive to the disruption of photomorphogenic development produced by retrograde signals. Finally, genetic interactions tested in *gun1-X* and *GLK1OX-X* mutants where endogenous *X* is removed, validate the involvement of X in the pathway.

(a)

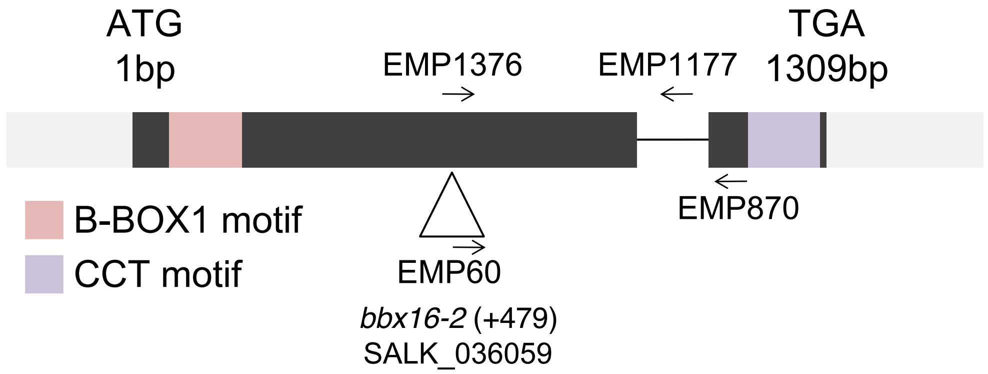


(b)

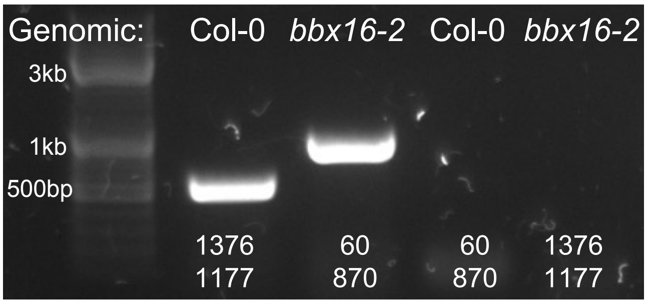


| | Primers: | Size: |
|---|---|---|
| Col-0 | EMP1376<br>EMP1177 | 530pb |
| *bbx16-2* | EMP60<br>EMP870 | ~1kb |

**Fig. S2. Molecular characterization of *bbx16-2*.**
**(a)** *bbx16-2* was obtained from the SALK collection (SALK_036059). It harbors a T-DNA insertion in the first exon, 479 base pairs downstream of the ATG. The position of the EMP primers used to genotype the line are indicated, as well as the the B-BOX1 and CCT motifs. **(b)** Agarose gel illustrating the genotyping of *bbx16-2* using Col-0 and *bbx16-2* genomic DNA and the indicated primer pairs to detect the WT band (EMP1376+EMP1177), and the mutant T-DNA band (EMP60+EMP870).

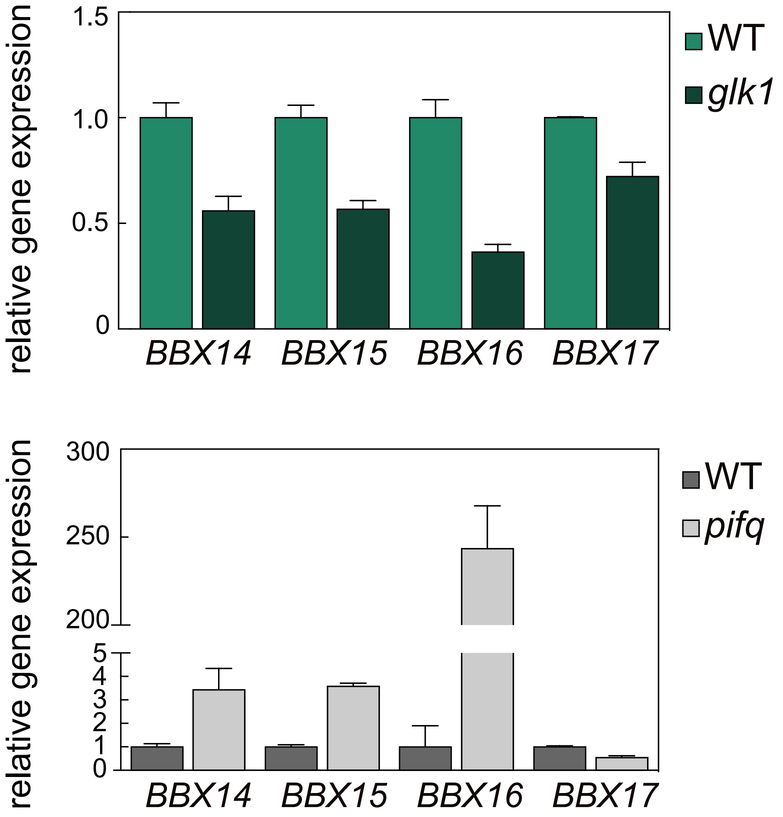


**Fig. S3. *BBX14*, *BBX15* and *BBX16* are similarly regulated by PIFs and GLK1 in dark and light.**

Expression of the four *BBXs* belonging to clade III of BBX family of transcription factors (*BBX14*, *15*, *16* and *17*) was analyzed by quantitative qRT-PCR in WT and *glk1* seedlings grown for 3 days in white light (5 $\mu mol \cdot m^{-2} \cdot s^{-1}$) (top) or from RNA-sequencing of WT and *pifq* seedlings grown for 3 days in dark (Martín et al., 2016) (bottom). Expression levels relative to wild type (WT) are shown. Data are the means ± SE of biological triplicates (except for *BBX17* expression value in the WT light samples (*n*=2)).

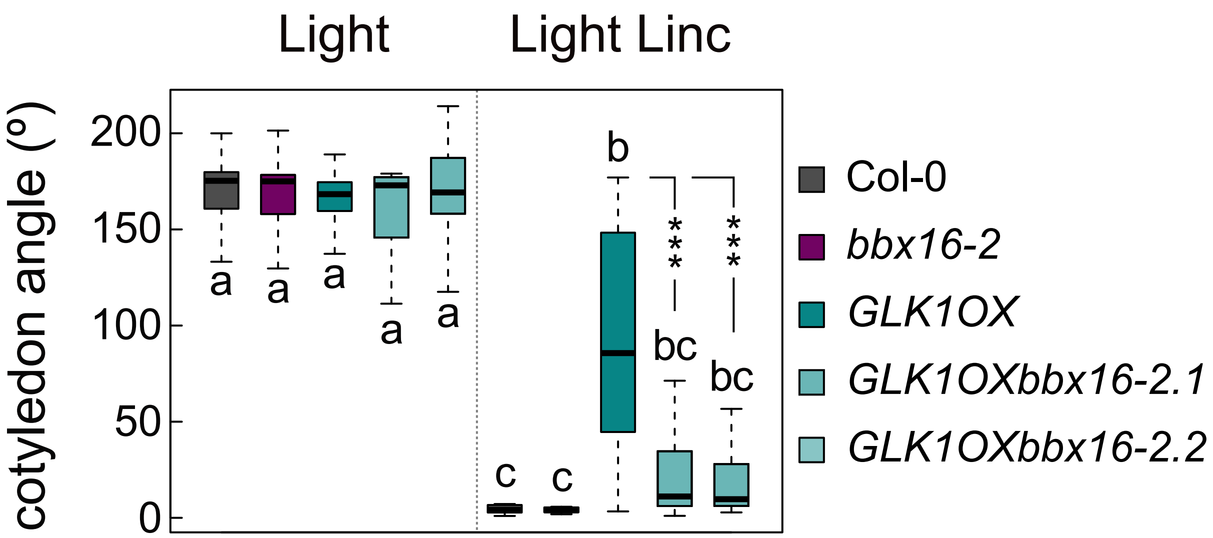


**Fig. S4. Genetic removal of *BBX16* partially suppresses the *GLK1OX* open cotyledon phenotype in the presence of lincomycin.**

Cotyledon angle of 3-day old light-grown (5 μmol·m$^{-2}$·s$^{-1}$) Col-0, *bbx16-2*, *GLK1OX*, and *GLK1OXbbx16-2* seedlings in the presence or absence of lincomycin. Data represent the median of at least 30 seedlings. Letters denote the statistically significant differences among genotypes by Kruskal-Wallis test ($P<0.05$), and asterisks indicate statistically significant differences between each *GLK1OXbbx16-2* mutant and *GLK1OX* seedlings (Mann-Whitney test; ***$P < 0.001$). Linc = Lincomycin.

**Supplemental Table 1: List of primers used for genotyping.**

Columns indicate the primer name, the sequence, and the genotyped allele. For primers described elsewhere, the reference is indicated.

| **Primer** | **Sequence** | **Allele** | **Reference** |
|---|---|---|---|
| EMP1472 | GCTTTCCTATGCCAAAGCTG | *bbx16-1* WT | |
| EMP870 | GTACCCACCCACAACATGAGG | *bbx16-1*, *bbx16-2* T-DNA | |
| EMP60 | TGGTTCACGTAGTGGGCCATCG | *bbx16-1*, *bbx16-2* T-DNA | |
| EMP1376 | CGGCTTTTGCTTAGTTCCTG | *bbx16-2* WT | |
| EMP1177 | AACGTCCGCAATGTGTTATTAAGTTGTC | *bbx16-2* WT | |
| EMP1091 | CCCAAAACAACACGAGTAGT | *glk1* WT | |
| EMP1096 | CACCTTTCCCCGAACTACCC | *glk1* WT | |
| EMP919 | CGGGATCCGACACTCTTTAATTAACTGACACTC | *glk1* T-DNA | (Fitter *et al.*, 2002) |
| EMP920 | AACTGCAGGTTACTGATCCGATTGTTCTT | *glk1* T-DNA | (Fitter *et al.*, 2002) |
| EMP1105 | CCACCAACACATATATACGGCA | *glk2* WT | |
| EMP1110 | TGTAACTCCGGCGTCCAATC | *glk2* WT | |
| EMP921 | CCTATTTCAGTAAGAGTGTGGGGTTTTGG | *glk2* T-DNA | (Fitter *et al.*, 2002) |
| EMP922 | CTCCTTGACGTTGTAACTCCG | *glk2* T-DNA | (Fitter *et al.*, 2002) |
| EMP884 | ATGCTGCATATCAGATTTCGG | *gun1* WT | |
| EMP883 | GTGGGTTCTGCTGTTTCTTG | *gun1* WT/T-DNA | |
| EMP92 | TAGCATCTGAATTTCATAACCAATCTCGATACAC | *gun1* T-DNA | |
| EMP1096 | CACCTTTCCCCGAACTACCC | *GLK1OX* | |
| 35S FWD | CGTTGAAGATGCCTCTGCCG | *GLK1OX* | |

**Supplemental Table 2: List of primers used for qRT-PCR.**

For each gene analyzed by qRT-PCR the pair of primer sequences is shown. Columns indicate the primer name, the sequence, and the gene amplified. For primers described elsewhere, the reference is indicated.

| **Primer** | **Sequence** | **Gene** | **Reference** |
|---|---|---|---|
| EMP338 | TATCGGATGACGATTCTTCGT | *PP2AA3* | (Shin *et al.*, 2007) |
| EMP339 | GCTTGGTCGACTATCGGAATG | *PP2AA3* | (Shin *et al.*, 2007) |
| EMP869 | TTGGGATAATCACGGTTCGCC | *BBX16* | |
| EMP870 | GTACCCACCCACAACATGAGG | *BBX16* | |
| EMP1180-P1 | GTAGGAGGACGCGCAGTAAT | *BBX16* | |
| EMP1182-P1 | AAAATGTTCCGTTGGTCGTC | *BBX16* | |
| EMP1175-P2 | TCATGCAAACGCACCAACAA | *BBX16* | |
| EMP1176-P2 | GTCATCTCCGTTTTTGCACTCT | *BBX16* | |
| EMP869-P3 | TTGGGATAATCACGGTTCGCC | *BBX16* | |
| EMP1177-P3 | GCTCAAAGTCAACGGAACTCG | *BBX16* | |
| PLR84 | CGTCCCCGGAAAGTGAGTT | *LHCB 1.4* | (Leivar *et al.*, 2008) |
| PLR85 | TGCAACAAACCGGATACACAC | *LHCB 1.4* | (Leivar *et al.*, 2008) |
| EMP945 | GCTTTGTAAACTCGTGATTGTG | *LHCB 2.2* | (Woodson *et al.*, 2013) |
| EMP946 | TGCCAAATTCACATCAAACG | *LHCB 2.2* | (Woodson *et al.*, 2013) |
| EMP1544 | GAAGGACTTGTGAAGGGAACA | *CA1* | (Hills *et al.*, 2015) |
| EMP1545 | TTTAACAGAGCTAGTTTCGGAGAG | *CA1* | (Hills *et al.*, 2015) |
| EMP1546 | AATTTCCGGACTTAACGTTTGTTT | *RBCS1A* | (Izumi *et al.*, 2012) |
| EMP1547 | CATCAGACAGTTGAGAATCCGATAGA | *RBCS1A* | (Izumi *et al.*, 2012) |
| EMP1548 | CCTATTGTCTGTGTTCTTTTTCTCTTTATG | *RBCS3B* | (Izumi *et al.*, 2012) |
| EMP1549 | TCAAGACGCACGGATATATAAATTACA | *RBCS3B* | (Izumi *et al.*, 2012) |
